# Genome editing of key domestication genes overcomes self-incompatibility and bitter taste in cultivated buckwheat

**DOI:** 10.64898/2026.05.27.728157

**Authors:** Artur Pinski, Magdalena Zaranek, Joanna Lusinska, Przemyslaw Kopec, Agnieszka Plazek, Paulina Pajak, Przemyslaw Petryszak, Anna Kostecka-Gugala, Meiliang Zhou, Alexander Betekhtin

## Abstract

Buckwheat (*Fagopyrum* spp.) is a climate-resilient pseudocereal, yet its global adoption is constrained by the distylous self-incompatibility of common buckwheat (*F. esculentum*) and grain bitterness of Tartary buckwheat (*F. tataricum*). While the *S-locus early flowering 3* (*FeS-ELF3*) gene has been identified as a key regulator of self-compatibility, a stable genetic transformation of *F. esculentum* has not yet been developed. In this study, we developed an Agrobacterium-mediated transformation protocol of *F. esculentum* (27% transformation efficiency) and applied it to the agronomically relevant ’Panda’ cultivar. Inactivation of the *FeS-ELF3* gene using the CRISPR/Cas9 system yielded self-compatible lines with long-homostylous flowers. *fes-elf3* mutants showed a distinct architectural shift: mutant plants were shorter and had shorter inflorescences than wild-type plants. Notably, these traits did not compromise yield, as the mutants produced a similar number of seeds per plant in the greenhouse. In *F. tataricum*, we targeted *the rutin-degrading enzyme (FtRDE2*), which was suspected to be a driver of grain bitterness by hydrolysing rutin into the bitter quercetin. Metabolic profiling of seeds of two *ftrde2* mutant lines revealed significantly lower quercetin levels in both. Analysis of enzyme extracts confirmed the loss of rutinosidase activity; the mutant samples maintained stable rutin concentrations without the characteristic increase in quercetin observed in the control. Furthermore, organoleptic sensory evaluation of flour demonstrated that respondents identified the control as significantly more bitter than the flour from the *ftrde2* mutants. These precise edits show proof-of-concept of overcoming domestication barriers: self-incompatibility and palatability, establishing a framework for rapid improvement of buckwheat.

## 1. Introduction

Buckwheat (*Fagopyrum* spp.), a member of the Polygonaceae family, comprises 26 recognised species, among which *Fagopyrum esculentum* Moench (commonly referred to as sweet or common buckwheat) and *Fagopyrum tataricum* (L.) Gaertn. (known as bitter or Tartary buckwheat) are the most widely cultivated. As an underutilised orphan crop, buckwheat has considerable potential to enhance global food and nutritional security. Naturally gluten-free and rich in high-quality proteins, essential minerals, and bioactive compounds such as rutin, buckwheat is considered a functional food. Its seeds are typically dehulled and milled into flour, which serves as a base for food products such as bread, noodles, pasta, cakes, biscuits, and fermented products such as wine, beer, and vinegar. The flowers are highly valued for honey production due to their distinctive colour and abundant nectar. Importantly, buckwheat is a resilient crop that can thrive in low-fertility, acidic soils and high altitudes, where many staple cereals fail. This adaptability makes it particularly suitable for cultivation in marginal environments and is relevant to sustainable agriculture (Ding *et al*., 2025; Jha *et al*., 2024; Pinski *et al*., 2025). Despite its nutritional and ecological value, sweet buckwheat suffers from self-incompatibility and relies on insect pollinators for fertilisation, leading to unstable yields (Kasajima *et al*., 2017). In contrast, Tartary buckwheat is self-compatible but produces seeds with an off-putting bitter taste(He *et al*., 2023; Yasui *et al*., 2012). To address these limitations, we developed genome editing tools targeting key genes identified through recent advances in buckwheat biology.

Successful fertilisation in *F. esculentum* occurs only between plants bearing complementary floral morphs—Thrum and Pin—due to a heteromorphic self-incompatibility (SI) system. This mechanism is genetically controlled by a hemizygous S-locus present in Thrum plants and is tightly linked to floral morphology. Previous studies have demonstrated that breakdown of SI and the emergence of long-homostyle flowers result from inactivation of *S-ELF3*, a homologue of *Arabidopsis thaliana ELF3*, located within the S-locus (Yasui *et al*., 2012). For instance, the *F. esculentum* PL4 line carries a nonsense mutation in *S-ELF3*, inherited from *F. homotropicum*. Additionally, ethyl methanesulfonate mutagenesis has produced a long-homostyle *F. esculentum* line due to splice site mutations in *S-ELF3* (Fawcett *et al*., 2023). This work relies on the analysis of single mutant alleles to establish gene–trait relationships at the S-locus. In the same study *F. tataricum*, a homomorphic and self-compatible species, structural variation in *S-ELF3* has been identified: an inverted duplication of the 5′ region and retrotransposon insertion. These findings from Fawcett *et al*. (2023) firmly establish *FeS-ELF3* as the key genetic determinant of SI in buckwheat.

Seed bitterness in *F. tataricum* is primarily caused by the activity of rutin-degrading enzymes (RDE), which catalyse the hydrolysis of rutin (quercetin 3-O-rutinoside) into quercetin, a bitter compound. In contrast, this issue is absent in *F. esculentum*, which has lower rutin content in seeds and lacks active RDE. Reducing bitterness is critical for enhancing consumer acceptance and marketability of *F. tataricum* products (Suzuki *et al*., 2014b; Wang *et al*., 2024; Zhao *et al*., 2024b; Zhao *et al*., 2025). RDEs in *F. tataricum* flour can be effectively inactivated through thermal treatments, including superheated and saturated steam. However, these methods are costly and can adversely affect flour properties, such as phenolic content and starch functionality (Kreft *et al*., 2022; Wu *et al*., 2020). To address this issue through breeding, the trace-rutinosidase line ‘f3g-162’—a Nepalese genetic resource—was used as the seed parent in a cross with ‘Hokkai T8’, a leading Japanese cultivar serving as the pollen parent. The resulting cultivar, ‘Manten-Kirari’, was found to exhibit only trace rutinosidase activity of ‘Hokkai T8’ and significantly reduced bitterness (Noda *et al*., 2022; Suzuki *et al*., 2014a). However, the specific gene(s) responsible for this phenotype were not fully understood. Early studies indicated that seed-expressed FtRDE comprises two distinct enzymes, F3gI and F3gII (also termed RDE I and RDE II). Subsequent research revised this view, identifying two or more isozyme variants—possibly differing in sugar composition—encoded by genes designated as Ft8.2377, FtBGLU29, or FtRDE2 (hereafter referred to as FtRDE2) (Jia *et al*., 2019; Noda *et al*., 2022; Suzuki *et al*., 2014a; Zhao *et al*., 2024b; Zhao *et al*., 2025). Specifically, expression analysis revealed that *FtRDE2* is highly expressed in *F. tataricum* seeds, with negligible or absent expression in roots, stems, leaves, or flowers. In contrast, its homolog in *F. esculentum* is not expressed due to multiple loss-of-function mutations (He *et al*., 2023). Further studies revealed that *FtRDE2* encodes a plasma membrane protein. Moreover, transient overexpression of *FtRDE2* in *F. tataricum* seeds significantly increased the rate of rutin hydrolysis (Zhao *et al*., 2025).

Thus, we established an Agrobacterium-mediated transformation protocol and targeted the *FeS-ELF3* gene to generate self-compatible *F. esculentum* plants. To improve the palatability of *F. tataricum*, we targeted the *FtRDE2* gene using a previously optimised transformation protocol developed by our group.

## 2. Results and Discussion

### 2.1 Inactivation of the *FeS-ELF3* gene leads to self-compatible *F. esculentum* plants

The sequencing of the coding sequence of *FeS-ELF3* confirmed the predicted intron-exon structure and revealed no single-nucleotide polymorphisms between *F. esculentum* “Pintian4” and *F. esculentum* cv. “Panda” (Supplementary Data 1). Given that *FeS-ELF3* is located within a hemizygous genomic region exclusive to Thrum plants (Fawcett *et al*., 2023), a Cas9 construct targeting *FeS-ELF3* was introduced into Thrum embryogenic calli induced from immature embryos using the Agrobacterium-mediated transformation system described here (Fig. S1a-b, Supplementary Data 1). The previously reported transformation protocol for *F. tataricum* was modified by the inclusion of a heat-shock step (43 °C for 5 min), which has been shown to increase transformation efficiency (Patel *et al*., 2013). This approach resulted in a transformation efficiency of 27.2% (6 out of 22 regenerated plants) and a genome-editing efficiency of 16.6% (1 out of 6 transgenic plants), which is comparable to the transformation efficiency (∼20%) achieved with our earlier optimised protocol for *F. tataricum* (Fig. S1c, Table 1). However, while transformation efficiency remained relatively high, the regeneration rate for *F. esculentum* is notably low; consequently, numerous transformation cycles were required to produce viable plants. Finally, targeted mutagenesis yielded a mutant carrying a single-nucleotide insertion in *FeS-ELF3*, resulting in a frameshift and the introduction of a premature stop codon, generating a loss-of-function allele (Fig. 1a, Fig S2a, Supplementary Data 1). The resulting *fes-elf3* mutant displayed long-homostylous flowers, a characteristic phenotype associated with the breakdown of self-incompatibility in *F. esculentum* (Fig. 1b). Reproductive potential was assessed by evaluating pollen viability in T0 plant. Although pollen viability in *fes-elf3* was slightly reduced relative to Thrum and Pin flowers, it remained within a functional range (Thrum, 88.9 ± 13.1%; Pin, 95.6 ± 4.1%; *fes-elf3*, 83.4 ± 10.1%; Fig. 1cd), indicating the capacity for successful fertilisation.

**Figure 1.**
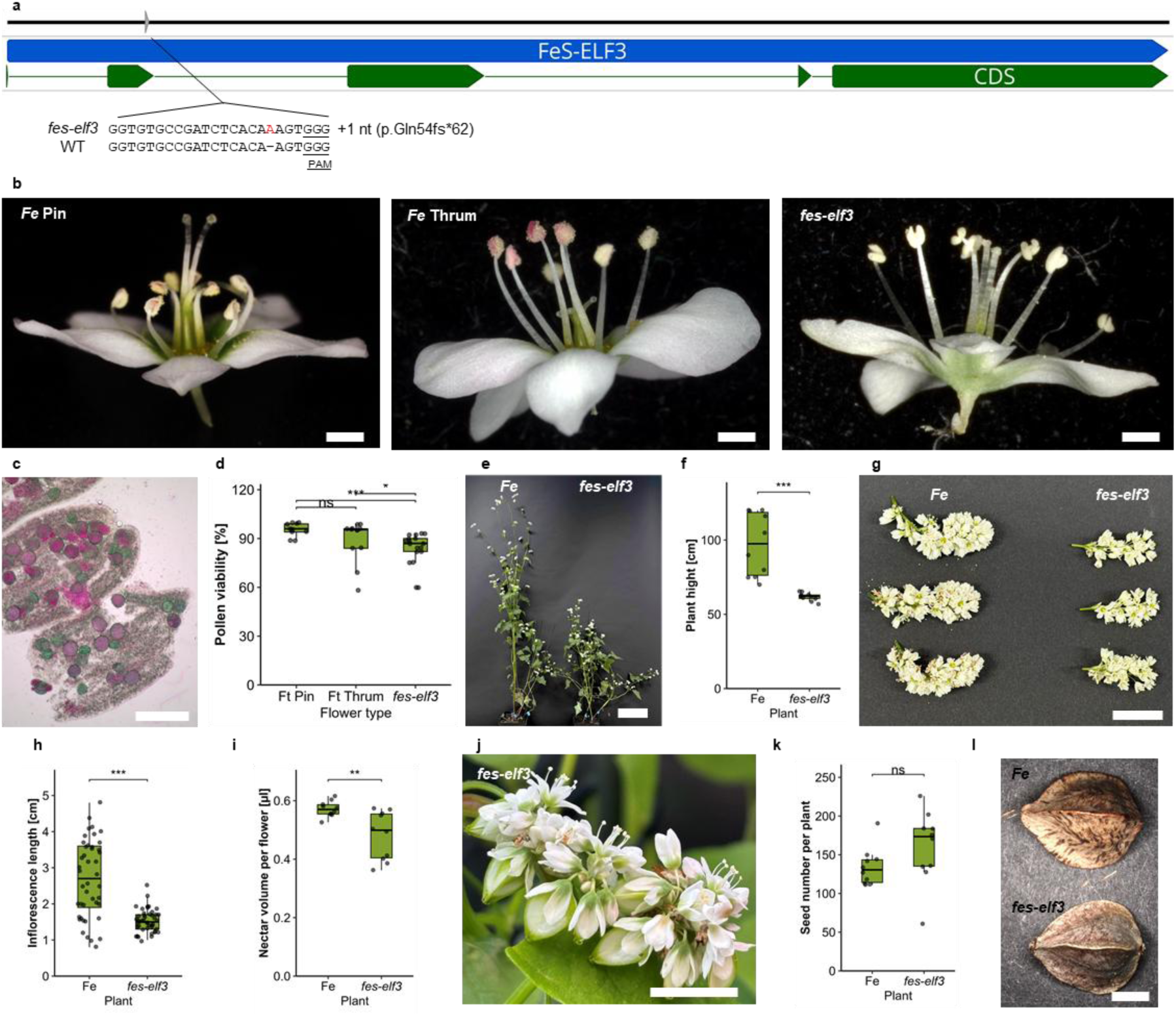
Inactivation of *FeS-ELF3* leads to long-homostylous flowers and self-compatibility. **a**, Schematic representation of the *FeS-ELF3* gene located in a hemizygous genomic region specific to Thrum plants, with the sgRNA target site indicated. PAM, protospacer adjacent motif. **b**, Floral morphotypes of *F. esculentum* cv. Panda. Pin flowers exhibit long styles and low anthers, whereas Thrum flowers have short styles and elevated anthers. Long-homostylous flowers result from nonsense mutations in *FeS-ELF3* from the T0 generation. Scale bar, 1 mm. **c**, Pollen viability of *fes-elf3* anthers from the T0 generation assessed by Alexander’s staining (Pin: n = 10; Thrum: n = 12; *fes-elf3*: n = 18). Scale bar, 100 µm. **d**, Quantification of pollen viability in Pin, Thrum, and *fes-elf3* (T0 generation) flowers. **e**, Representative photographs of *fes-elf3* (T2 generation) and WT plants grown under greenhouse conditions. Scale bar, 10 cm. **f**, Plant height of *fes-elf3* (T2 generation) and WT plants grown in the greenhouse (n = 10 plants per genotype). **g**, Representative inflorescences of *fes-elf3* (T2 generation) and WT plants. Scale bar, 2 mm. **h**, Inflorescence length of *fes-elf3* (T2 generation) and WT plants (n = 9 plants per genotype; 5 inflorescences per plant). **i**, Nectar volume per flower in *fes-elf3* (T2 generation) and WT plants (n = 10 flowers per genotype). **j**, Inflorescence of *fes-elf3* (T2 generation) showing flowers and developing seeds. Scale bar, 1 cm. **k**, Seed number per plant in *fes-elf3* (T2 generation) and WT under greenhouse conditions (n = 10 plants per genotype). **l**, Representative seeds of *fes-elf3* (T2 generation) and WT plants. Scale bar, 2 mm. Boxplots represent the median and the interquartile range of the data. Statistical significance was determined by Student’s *t*-test (**f**,**h, i, k**) or Dunn’s test following Kruskal–Wallis (**d**). Significance levels are indicated as ***P < 0.001; **P < 0.01; ns, not significant.

**Table 1:** Summary of Agrobacterium-mediated transformation and CRISPR/Cas9 editing efficiency in *Fagopyrum* species. Mutation frequency is calculated as the percentage of T-DNA-integrated plants exhibiting targeted sequence modifications.

| Species | Target Gene | Regenerated Plants (T0) | Transgenic Plants (T-DNA+) | Edited plants (mutation frequency) |
| --- | --- | --- | --- | --- |
| *F. esculentum* | *FeS-ELF3* | 22 | 6 (27.27%) | 1 (16.67%) |
| *F. tataricum* | *FtRDE2* | 83 | 19 (22.89%) | 13 (68.42%) |

T1 progeny were resequenced to confirm the inheritance of the targeted *FeS-ELF3* mutations identified in the parental T0 line (Fig. S2b). Subsequently, transgene-free T2 individuals were identified and characterised under greenhouse conditions. This stable transmission of edited alleles to the T1 generation demonstrates the effectiveness of this strategy for developing new buckwheat varieties. We observed that *fes-elf3* plants were significantly shorter than WT plants, with average heights of 61.9 ± 2.7 cm compared with 97.0 ± 21.1 cm in WT (Fig. 1e, f). In addition to reduced plant height, *fes-elf3* plants exhibited shorter inflorescences (1.5 ± 0.3 cm) relative to WT plants (2.7 ± 1.1 cm; Fig. 1g, h). Importantly, subsequent analysis of five consecutive generations (T0-T4) revealed no changes in plant phenotype, confirming the stable, heritable nature of the edited trait across multiple generations. This more compact growth habit may be advantageous for future agricultural applications, as lodging is a major constraint on grain yield and the development of the buckwheat industry (Hornyák *et al*., 2020; Xiang *et al*., 2019). Indeed, a previously developed ethyl methanesulfonate–induced dwarf mutant of *F. tataricum* (*ftdm*) was shown to be suitable for high-density planting and commercial cultivation (Sun *et al*., 2021). Previously, differences in plant height or inflorescence length in *fes-elf3* mutants had not been reported, as *FeS-ELF3* is primarily known to regulate style length and female compatibility (Fawcett *et al*., 2023; He *et al*., 2023; Lin *et al*., 2023; Ma *et al*., 2022). However, given its role as a transcription regulator, it is plausible that *FeS-ELF3* also regulates additional downstream genes involved in vegetative growth and inflorescence development (Yasui *et al*., 2012), necessitating further analysis. Because *F. esculentum* is an important nectar source for pollinators (Plazek *et al*., 2023), nectar production was also evaluated. Nectar volume per flower was moderately reduced in *fes-elf3* plants (0.481 ± 0.083 µL) compared with WT plants (0.572 ± 0.027 µL; Fig. 1i). *fes-elf3* plants maintained in a separate growth chamber was able to set seeds in the absence of cross-pollination, providing direct evidence of self-compatibility (Fig. 1j). Seed number per plant was comparable between WT (134.2 ± 24.6) and *fes-elf3* (160.3 ± 46.8) plants under greenhouse conditions (Fig. 1k, l). A large-scale field experiment will be necessary to accurately assess the yield of *fes-elf3* plants. However, it is expected that these self-compatible plants will provide more stable year-to-year yields, as self-fertilising plants are independent of pollinator availability (Kasajima *et al*., 2017). In contrast, pollination success in self-incompatible plants depends on unpredictable weather events, an effect that is further exacerbated by the short one-day pollination window of each flower (Cawoy *et al*., 2009).

### 2.2. Inactivation of *FtRDE2* genes leads to non-bitter seeds of *F. tataricum* plants

Using a previously established Agrobacterium-mediated transformation protocol for *F. tataricum* (Pinski and Betekhtin, 2023), we obtained 83 regenerated T0 plants, of which 19 were confirmed as transgenic (22.89% T-DNA integration efficiency) (Table 1). Within this transgenic population, we identified 13 independent *ftrde2* mutants, representing a mutation frequency of 68.42%. Sequence analysis revealed that CRISPR/Cas9-mediated editing predominantly induced small deletions, ranging from –1 to –6 bp, although larger deletions (up to –98 bp) and small insertions (+1 to +4 bp) were also observed (Fig. S3). For further phenotypic analysis, we selected two independent mutant lines: *ftrde2_1*, characterised by a clear frameshift, and *ftrde2_2*, which had small in-frame deletions. In *ftrde2_1*, biallelic insertions of 1 and 40 nucleotides disrupted the reading frame, introducing premature stop codons and resulting in a complete loss-of-function. In contrast, *ftrde2_2* harboured deletions of 3 and 6 nucleotides, resulting in the loss of 1 and 2 amino acids, respectively (Fig. 2a). Despite being in-frame, these modifications successfully inactivated the enzyme, as confirmed by subsequent phenotypic characterisation (Fig. 2b-f). In our previous study targeting the *FtPDS* gene in *F. tataricum* (Pinski and Betekhtin, 2023), we observed primarily small indels induced. In contrast, the *ftrde2_1* line characterised here contains a 40-nt insertion at the target site. BLAST analysis of this sequence against the *F. tataricum* genome revealed high similarity to the intron located immediately downstream of the target sequence. This 40-nt insertion shares 76.2% identity with the intronic region, with a central 20-nucleotide stretch identical to the genomic reference, suggesting a templated insertion event during double-strand break repair (Fig. S4). We subsequently obtained T1 progeny from the transgenic lines and confirmed their genotypes via sequencing (Fig. S5, Supplementary Data 1). The seeds harvested from these transgene-free verified T1 individuals were utilised for all downstream phenotypic and biochemical analyses.

**Figure 2:**
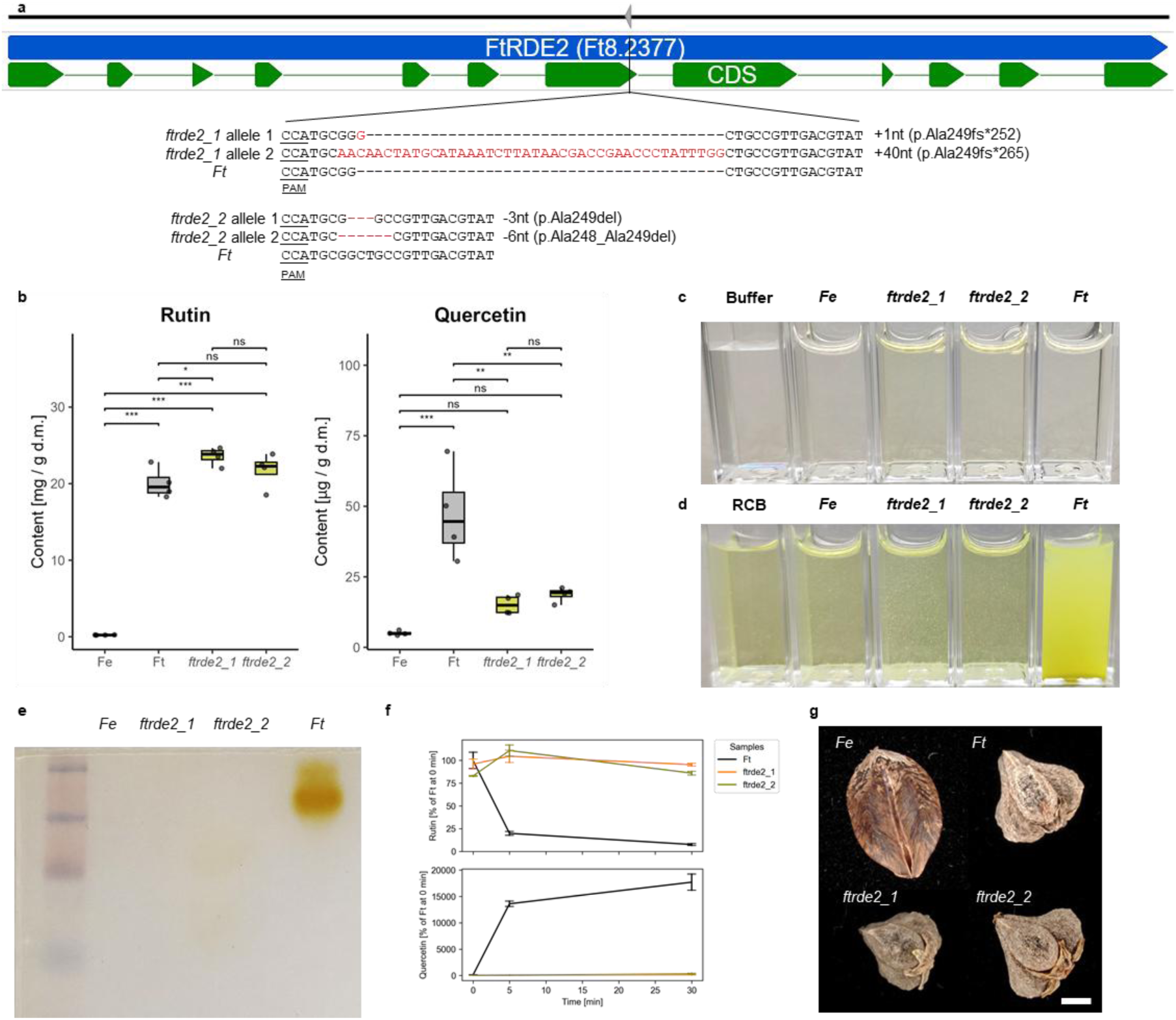
Inactivation of an *FtRDE2* results in non-bitter seeds. **a**, Schematic representation of the *FtRDE2* gene, which encodes an RDE, with the sgRNA target site indicated. Sequence results are shown for two independent T0 mutant lines (*ftrde2_1* and *ftrde2_2*) selected for further phenotypic characterisation. PAM – protospacer adjacent motif. **b**, Rutin and quercetin concentration in seed flour from F. *esculentum*, *F. tataricum*, f*trde2_1*, and *ftrde2_2* (n = 4 biological replicates per genotype). **c**, Crude enzyme extracts prepared from seed flour of *F. esculentum*, *ftrde2_1*, *ftrde2_2*, and *F. tataricum*. **d**, Enzymatic activity of crude seed flour extracts from *F. esculentum*, *ftrde2_1*, *ftrde2_2*, and *F. tataricum* analysed in rutin-containing buffer (RCB). **e**, Detection of rutin-degrading enzyme activity in *F. esculentum*, *ftrde2_1*, *ftrde2_2*, and *F. tataricum* using native PAGE. **f**, Rutinosidase activity assay using crude seed flour extracts from *ftrde2_1*, *ftrde2_2*, and *F. tataricum*, showing changes in rutin and quercetin levels at 0, 5, and 30 min (n = 3 biological replicates per genotype). **g**, Photographs of seeds from *F. esculentum*, *ftrde2_1*, *ftrde2_2*, and *F. tataricum*. Scale bar, 1 mm. Boxplots represent the median and the interquartile range of the data. Statistical significance was determined by Dunn’s test following Kruskal–Wallis (**b**). Significance levels are indicated as ***P < 0.001; **P < 0.01; ns, not significant.

Measurement of rutin content in seed flour revealed no significant difference between WT (*F. tataricum*, 20.06 ± 2 mg/g dry mass) and *ftrde2_2* (21.73 ± 2.27 mg/g dry mass) (Fig. 2b). While *ftrde2_1* exhibited slightly higher rutin content (23.59 ± 1.16 mg/g dry mass) compared to WT, all *F. tataricum* maintained significantly higher levels than *F. esculentum* flour (0.22 ± 0.02 mg/g dry mass). These values are consistent with previous studies demonstrating the high rutin content of *F. tataricum*, comparable to that of the superior cultivar Donan (21.31 mg/g dry mass; Brunori *et al*. (2007)). Measurement of quercetin content revealed 15.17 ± 3.32 µg/g dry mass in *ftrde2_1* and 18.7 ± 2.6 µg/g dry mass in *ftrde2_2*, which were not significantly different from *F. esculentum* (5.04 ± 0.75 µg/g dry mass), but substantially lower than in *F. tataricum* (47.32 ± 16.8 µg/g dry mass). In agreement with previous reports, quercetin was present at much lower levels than rutin in buckwheat plant tissues. Unlike rutin, which generally increased during plant growth, quercetin levels in *F. tataricum* flowers gradually declined over the same period (Bai *et al*., 2015; Kim *et al*., 2023; Zielinska *et al*., 2012). Consistent with the rutin and quercetin measurements, crude seed enzyme extracts from *ftrde2_1* and *ftrde2_2* displayed a yellow coloration characteristic of the flavonol chromophore (Abualhasan *et al*., 2017), absent in WT *F. tataricum* — where rutin is rapidly degraded — and in *F. esculentum*, where seed rutin concentration is approximately 90-fold lower than in *F. tataricum* (Fig. 2c, d). When degradation of exogenous rutin was tested using crude enzyme extracts, no rutin degradation was observed in *F. esculentum* or in the *ftrde2_1* and *ftrde2_2* mutants, whereas the wild-type *F. tataricum* extract efficiently degraded rutin into quercetin and rutinose, resulting in increased turbidity. *F. esculentum* served as a negative control, as its seeds lack RDE activity (Fig. 2d; He *et al*. (2023)). Furthermore, native PAGE assays confirmed the absence of rutin-degrading enzyme activity in both mutants. No activity was detected in *F. esculentum*, *ftrde2_1*, or *ftrde2_2*, whereas a distinct activity band was observed in wild-type *F. tataricum*, indicating the presence of a functional RDE (Fig. 2e). We also investigated rutinase activity in crude seed flour extracts at 0, 5, and 30 min in a standard mixture that contains exogenous rutin (0.2% w/v) (Fig. 2f). In wild-type *F. tataricum*, most rutin was degraded within 10 min, coinciding with an increase in quercetin. In contrast, rutin levels in the *ftrde2_1* and *ftrde2_2* mutants remained constant over time, and quercetin levels also showed no change, confirming the complete loss of RDE activity in these mutants (Fig. 2f). These results are consistent with previous reports on the non-bitter, trace-rutinosidase variety of *F. tataricum*, ‘Manten-Kirari’ (Suzuki *et al*., 2014a).

We further assessed the bitterness of flour from the *ftrde2* mutant seeds using sensory evaluations. Organoleptic testing of flour prepared from *F. tataricum* seeds revealed that in paired comparison tests, 43 out of 45 (95.5%) and 42 out of 45 (93.3%) respondents correctly identified the control flour as more bitter than that from *ftrde2_1* and *ftrde2_2*, respectively (Supplementary Data 2). These results were highly significant (one-tailed binomial test, *p* < 0.001), confirming a clear reduction in perceived bitterness. Moreover, the inactivation of the gene did not visibly affect seed morphology compared to the WT (Fig. 2g). Our results clearly demonstrate that *FtRDE2* is the primary determinant of seed bitterness in *F. tataricum*, and its inactivation produces non-bitter seeds with high rutin content, as previously hypothesised (He *et al*., 2023; Wang *et al*., 2024; Zhao *et al*., 2024a; Zhao *et al*., 2025).

While the observed phenotype of *ftrde2_1* can be readily explained by frameshift mutations in both alleles leading to a premature stop codon, the in-frame mutations in *ftrde2_2* were surprising in their ability to cause the observed RDE inactivation. The deletion of a single ALA-218 or double ALA-218-219 in *ftrde2_2* alleles does not occur in the proximity of the previously described rutin-binding pocket (positions are numbered after removal of the signal peptide, residues 1–30 as done in Zhao *et al*. (2025)). It was shown that rutin has a strong affinity for FtRDE2, fitting into a molecular pocket formed by LEU-178, ASN-311, GLU-383, LEU-356, and THR-243. Furthermore, rutin forms stable hydrogen bonds with GLU-383 and SER-355 and engages in π–π stacking interactions with TRP-434 (Fig S6). In our AutoDock Vina analysis, the ΔAla218 and ΔAla218–219 variants showed slightly more favourable predicted binding energies than WT (−8.871 and −8.767 vs −8.499 kcal/mol; Supplementary Data 3). However, because docking scores primarily reflect binding energetics rather than catalytic competence, these predictions are best interpreted as consistent with the possibility that the in-frame deletions disrupt productive substrate positioning, local dynamics, or coupling between binding and catalysis, rather than simply weakening ligand affinity.

In summary, our study demonstrates the feasibility and effectiveness of genome editing in buckwheat as a strategy to overcome long-standing breeding challenges. We present a transformation protocol for *F. esculentum*, in addition to our previously published method for *F. tataricum*. Through precise genome editing, we address two major limitations in buckwheat cultivation: self-incompatibility in *F. esculentum* and seed bitterness in *F. tataricum* caused by rutin-degrading enzymes. Notably, we confirmed the stable transmission of these targeted modifications to the T1 generation, establishing that these edited traits are heritable. The obtained mutants were generated using the CRISPR/Cas9 system, a new genomic technique (NGT) recognised for its precision in genome editing. Although opinions differ globally on whether CRISPR-derived crops should be classified as GMOs, such plants are generally more widely accepted and face fewer regulatory restrictions, allowing faster adoption in agriculture (Menary and Fuller, 2024). *F. esculentum* cv. Panda is a commercially available variety grown in Poland; therefore, the self-compatible line could be cultivated directly or introduced into other elite breeding lines (Suzuki *et al*., 2014a). Similarly, *rde2* mutants of *F. tataricum* can serve directly as a functional food due to its rich nutritional profile or as valuable material for breeding programs (Mukasa *et al*., 2007). Ultimately, this approach enables the development of elite cultivars with stacked traits, including dwarfism, waxy starch, reduced allergenicity, and enhanced tolerance to abiotic stress, thereby contributing to resilient and sustainable food systems.

## 3. Experimental procedures

### Plant material

Morphogenic calli of *F. tataricum* and embryogenic calli of *F. esculentum*, both derived from immature zygotic embryos, were used for Agrobacterium-mediated transformation. In the case of *F. esculentum*, calli were induced from Thrum immature zygotic embryos. Once sufficient tissue was obtained, genomic DNA was isolated using the CTAB method. A fragment of the *FeS-ELF3* gene was subsequently amplified (FeS-ELF3_partial_F and FeS-ELF3_partial_R) using Q5 High-Fidelity DNA Polymerase (NEB, UK) and sequenced to verify the Thrum-type origin of the callus and confirm the presence of the *FeS-ELF3* gene (Fig. S1a, Supplementary Data 1). The oligonucleotides used in this study are listed in Supplementary Data 4. The callus was maintained on RX medium according to (Betekhtin *et al*., 2017) at 25±1°C in the dark and subcultured every three to four weeks for *F. tataricum* and every two weeks for *F. esculentum*. Before transformation, embryogenic Thrum type callus of *F. esculentum* was put on regeneration medium containing macro- and micro-elements as in MS medium (Murashige and Skoog, 1962), 3.0 mg/L 6-benzylaminopurine, 1.0 mg/L thidiazuron, 30 g/L sucrose, 3 g/L phytagel and cultured in a growth room at 28 ± 2°C with a 16/8 h (light/dark) photoperiod, under light intensity of 55 µmol/m^2^/s for 12-14 days to induce somatic embryos. Callus with forming somatic embryos was transformed.

### Cloning of *FeS-ELF3* and *FtRDE2* genes, vector design, and selection of CRISPR/Cas9 target sites

To validate the sequence and exon-intron structure of the *FeS-ELF3* gene in *F. esculentum* cv. Panda, total RNA was isolated from young Thrum-type flowers using the RNeasy Mini Kit (Qiagen, Hilden, Germany). First-strand cDNA was synthesised using the Maxima H Minus First Strand cDNA Synthesis Kit with dsDNase (Thermo Scientific, Waltham, MA, USA). The complete *FeS-ELF3* coding sequence was subsequently amplified using Q5 High-Fidelity DNA Polymerase (NEB, UK) with the primers BsaI_FeS-ELF3_F and BsaI_FeS-ELF3_R. These oligonucleotides were designed using the *F. esculentum* “Pintian4” reference genome from the Chinese National Genomics Data Center (GWHBJBK00000000). The oligonucleotides were designed to remove the STOP codon and add BsaI recognition overhangs, allowing Golden Gate cloning into vector pGGC000 from the GreenGate cloning system. The vectors with cloned *FeS-ELF3* were sequenced using M13F and SP6 oligonucleotides (Supplementary Data 1). Likewise, a fragment of the FtRDE2 gene was amplified using FtRDE2_partial_F and FtRDE2_partial_R oligonucleotides and cloned into the pGEM-T Easy vector. The oligonucleotides were designed using the *F. tataricum “Pinku1”* reference genome from the Chinese National Genomics Data Center (GWHBJBL00000000) (Supplementary Data 1).

For targeted mutagenesis, we used the binary vector pGG-Cas9-AtU6-lacZ-kanR, constructed using the GreenGate cloning system (Lampropoulos *et al*., 2013). The empty vector was assembled as described in Pinski and Betekhtin (2023) with two modifications: pGG-Cas9(ΔSTOP) was replaced with pGG-Cas9(STOP), and the pGGD001 (linker-GFP) module was substituted with pGGD002 (C-tag, D-dummy) (Fig. S7, GeneBank accession number: PZ366425). The gRNA sequences targeting the *FtRDE2* and *FeS-*

*ELF3* genes were designed using Geneious Prime 2022.0.2, with off-target site predictions performed against the *F. tataricum* “Pinku1” and *F. esculentum* “Pintian4” genomes, respectively. Off-target sites were assessed against the reference genome assembly using Geneious specificity scoring, allowing up to 3 mismatches and no indels. Final sgRNAs were selected based on high predicted activity, proximity to the 5′ coding region, and the absence of predicted off-target sites within the reference genome under these criteria. Target-specific sgRNA sequences were introduced into the vector pGG-Cas9-AtU6-lacZ-kanR via Golden Gate assembly using flanking AarI restriction sites, and the assembly was confirmed by sequencing. Assembled vectors were electroporated into the *A. tumefaciens* GV3101 strain.

### Agrobacterium-mediated transformation

Agrobacterium preparation, callus transformation, and plant regeneration were performed according to the protocol previously described by Pinski and Betekhtin (2023) for Agrobacterium transformation of *F. tataricum*, with minor modifications. For *F. esculentum,* Thrum type callus with somatic embryos (5-7 g) was placed in 30 mL of liquid RX medium supplemented with 30 mg/L acetosyringone and 0.1% Pluronic F-68 for heat shock in a water bath at 43 °C for 5 min. Afterwards, the RX liquid medium was removed, and the tissue was incubated in 30 mL of Agrobacterium suspension for 10 min. Subsequently, the Agrobacterium liquid was removed, and the tissue was dried on 110 mm filter papers in open Petri dishes for 15 min. The next steps were the same as for *F. tataricum* Agro-transformation. To minimise the occurrence of chimeric plants, the kanamycin concentration used for selection was increased from 50 mg/L to 100 mg/L, and the selection period was extended from 3–4 weeks to 6–7 weeks.

### Regenerated plants cultivation

Regenerated plants with developed roots were transferred to an aero-hydroponic propagator for 20 cuttings (Hemp) filled with medium containing 20 ml of Aqua Vega A (Canna, Netherlands) and 20 ml of Aqua Vega B (Canna, Netherlands) per 10 L of water. For the first week, the lid was applied to maintain humidity; after this time, the lid was gradually removed to acclimate the plants (Fig. S1b). Rooted and strong plants were transferred to pots filled with moss-coconut fibre substrate (Ceres International) and grown under greenhouse conditions (18–26 °C, 16/8 photoperiod, light intensity of 40 µmol/m^2^/s sodium lamps Lucalox LU600W/PSL, HU, optimally irrigated). Seeds from the T0 generation were collected in successive rounds. The *fes-elf3* plants were maintained in a greenhouse chamber without any other *F. esculentum* plants to prevent cross-pollination. The nectar analysis was performed as described previously (Plazek *et al*., 2023). Wild-type *F. esculentum* plants were grown in a separate chamber containing *Calliphora vicina* flies, enabling efficient cross-pollination between Pin and Thrum plants.

### Mutants characterisation

The DNA from the leaves of regenerated plants was isolated using the CTAB method, as previously described by Hus et al. (Hus *et al*., 2020). Successful transformation and T-DNA integration were first confirmed by PCR amplification of a fragment of the Cas9 gene (Cas9_partial_F and Cas9_partial_R). For successfully transformed *F. tataricum* plants, a partial sequence of the *FtRDE2* gene was amplified and cloned into the pGEM-T Easy vector. In the case of *FeS-ELF3*, because a single copy of the gene is present in *F. esculentum* Thrum plants, the PCR product was directly sequenced. The resulting sequencing data were aligned with the corresponding reference sequences using Geneious Prime 2022.0.2 to identify potential mutations within the target genes and to characterise potential influence on protein formation. The Sanger sequencing results for WT and mutant sequences are in Supplementary Data 1. The transgene-free plants were assessed by PCR amplification of a fragment of the Cas9 gene (Cas9_partial_F and Cas9_partial_R).

### Microscopic observations

The bright field images of the flower and seeds were captured using a KEYENCE VHX-7000 digital microscope (Keyence, Japan).

### Native polyacrylamide gel electrophoresis (PAGE)

Seeds of *F. esculentum*, *F. tataricum* WT, and the *ftrde2_1* and *ftrde2_2* mutants (seeds collected from T1 transgene-free plants) were ground and sieved to obtain flour. Fifty milligrams of flour were extracted with a buffer containing 10% (v/v) glycerol and 0.01% (w/v) Triton X-100 in 50 mM sodium acetate–NaOH buffer (pH 5.0). The resulting crude enzyme extract was subjected to 7.5% (w/v) native-PAGE under non-denaturing conditions using a slab gel apparatus (Bio-Rad). Following electrophoresis, the gel was equilibrated in 50 mM sodium acetate–NaOH buffer (pH 5.0, 4°C) containing 20% (v/v) methanol for 10 min. The equilibrated gel was then incubated for 20 min in rutin-containing buffer (50 mM sodium acetate–NaOH buffer, pH 5.0, 4°C, 20% (v/v) methanol, 0.6% (w/v) rutin, and 5 mM CuSO₄) as described by Suzuki *et al*. (2004), and subsequently washed thoroughly with water. For visual assessment of rutinosidase activity, 50 µL of crude enzyme extract from each sample (*F. esculentum*, *F. tataricum* WT, *ftrde2_1*, and *ftrde2_2*) was added to 1 mL of rutin-containing buffer, and the mixtures were incubated for 10 min before making photographs.

### Organoleptic analysis

For this assay, seeds collected from T1 transgene-free plants of *ftrde2_1* and *ftrde2_2* were used. A panel of 15 assessors from the Faculty of Food Technology (University of Agriculture in Krakow) was selected for the test based on their experience in sensory evaluation. The panellists had been regularly trained and tested in accordance with ISO sensory standards (e.g. ISO 5492:2008, ISO 8586:2023, ISO 8587:2006, ISO 5495:2005, ISO 4120:2021, ISO 10399:2017, ISO 11132:2021, ISO 3972:2011). The analysis was conducted in individual booths with uniform lighting conditions at the Laboratory for Sensory Analysis (part of the Centre for Innovation and Research on Pro-Healthy and Safe Food, University of Agriculture in Krakow), designed in accordance with ISO 8589:2007. Buckwheat flour samples (WT and *ftrde2_1*) were served on identical plastic cups, labelled with random numbers, and arranged in a randomised order. The samples were assessed in pairs by panellists in three repetitions, yielding 45 individual results. Distilled water and apple slices were used as taste neutralisers between the samples. A second identical pair test was conducted in a subsequent session to compare the control sample with *ftrde2_2*. The research activities regarding sensory analysis were approved by the University of Agriculture Ethics Committee (Approval No. 283/2025, June 23, 2025), and all participants provided informed consent. The results of the pair comparison test were analysed using a one-tailed binomial test (H₀: P = 0.5; H₁: P > 0.5), where *p* represents the probability of correctly identifying the more intense (bitter) sample. The binomial test was applied to determine whether the observed proportion of correct responses differed significantly from the chance level. Statistical significance was defined as *P* < 0.05. All analyses were performed using Statistica.

### Pollen viability

Pollen viability (T0 generation of *fes-elf3* mutant) was assessed according to Alexander’s staining method (Alexander, 1969). Fully open flowers were collected from the greenhouse-grown plants. Then, fresh pollen was put on a drop of Alexander’s stain on microscope slides (samples were taken from 6 randomly chosen flowers) and covered with a cover glass. The slides were examined under an inverted Leica DMi8 microscope (Leica Microsystems, Germany) equipped with a Leica DFC 7000 T camera conjugated with LAS X Extended Depth of Field and Deconvolution Modules. The number of viable (dark red cytoplasm with a green wall) and non-viable (entirely green) pollen grains was counted. Pollen viability was expressed as the percentage of viable pollen.

### Rutin and quercetin extraction

For this assay, seeds collected from T1 transgene-free plants of *ftrde2_1* and *ftrde2_2* were used. To determine rutin and quercetin content in flour using high-performance liquid chromatography (HPLC), the extracts were prepared by vigorously vortexing about 20 mg of sample in 1 mL of 80% methanol for 30 min at room temperature. The homogenates were then centrifuged (15,000 × g, 10 min, 20°C). The resulting supernatants were filtered through a 0.45-micron nylon syringe filter (Millipore Millex-HN, Burlington, MA, USA). Four biological replications were measured.

### Rutinosidase activity analysis

For this assay, seeds collected from T1 transgene-free plants of *ftrde2_1* and *ftrde2_2* were used. The measurements were carried out in three biological replications. Rutinosidase activity was analysed by measuring quercetin and rutin concentrations in the reaction mixture using HPLC. The standard assay mixture consisted of 50 mM acetate–NaOH buffer (pH 5.0 at 4°C), 20% (v/v) methanol, and 0.2% (w/v) rutin (Sigma-Aldrich, Merck KGaA, Darmstadt, Germany) in a final volume of 0.05 mL (Suzuki *et al*., 2002; Suzuki *et al*., 2014a). The rutinosidase activity was expressed relative to the initial rutin concentration in *F. tataricum* WT reaction mixture. To prepare the reaction mixture, about 20 mg of Tartary buckwheat flour was thoroughly mixed with 0.2 mL of the standard assay mixture and stored at 37°C for 0, 5, and 30 min. 0.8 mL of methanol was added to the reaction mixtures to inactivate rutinosidase. The mixtures were centrifuged (15,000 × g, 10 min, 20°C) and the resulting supernatants were filtered through a 0.45-micron nylon syringe filter (Millipore Millex-HN, Burlington, MA, USA).

### Evaluation of rutin and quercetin content by HPLC

Amounts of rutin and quercetin were evaluated using UFLC Shimadzu Prominence liquid chromatography system (equipped with a SPD M20A Diode Array Detector, CTO-10AS VP Column Oven, and a SIL-20AC HT Autosampler; Shimadzu, Kyoto, Japan). A C18 RP chromatographic column (LiChrospher® 100 RP-18, 250–4, 5 µm, Merck KGaA, Darmstadt, Germany) was used. Chromatographic separations were performed at 30°C using the following solvents: (A) water with acetic acid (0.1%), (B) 50% methanol with acetic acid (0.1%), (C) 100% methanol with acetic acid (0.1%), and using the gradient: 100% A at the start, increasing to 100% B by 38 min, then to 100% C by 76 min, followed by a 4-min hold at 100% C. Total separation time, excluding column wash and equilibration, was 110 min. The flow rate was 1 mL/min. Qualitative analysis was performed by comparing the retention times of the analysed peaks and the peaks of analytical standards of rutin and quercetin (both from Sigma-Aldrich, Merck KGaA, Darmstadt, Germany) as well as by analysing their absorption spectra in the 190–400 nm range, as displayed by the chromatograph. Quantitative analysis was performed based on chromatograms obtained at 260 nm wavelength. The standards were dissolved in 80% methanol and stored at –20°C in the dark. The water and methanol used were HPLC-grade (Sigma-Aldrich, Chromasolv for HPLC), and the acetic acid was analytical grade.

### 3D structure modelling and molecular docking of FtRDE2 variants

Three-dimensional structures of *FtRDE2* wild-type, *ftrde2_2* ΔALA218, and *ftrde2_2* ΔALA218ΔALA219 were generated as follows (Fig. S6; Supplementary Data 3). The N-terminal signal peptide (residues 1–30) was removed from each sequence prior to structure prediction using AlphaFold3 (Abramson *et al*., 2024). The resulting predicted structures were then subjected to molecular docking with rutin using AutoDock Vina (https://www.molecular-modelling.ch/swiss-drug-design.html) to calculate binding free energies and generate multiple binding conformations (Bugnon *et al*., 2024; Eberhardt *et al*., 2021). Binding poses were ranked by calculated affinity (kcal/mol), and the top-scoring models were visualised using PyMOL. Previously predicted rutin-binding residues were annotated in accordance with a prior report (Zhao *et al*., 2025).

### Statistical analysis

All statistical analyses and data visualisations were performed in RStudio (R version 4.4.1; RStudio PBC, Boston, MA, USA) using the following R packages: readxl, dplyr, tidyr, purrr, rstatix, ggplot2, ggsignif, patchwork, and RColorBrewer.

## Authors contributions

Conceptualization: AP, AB; Methodology: AP, MZ, JL, PK, AG, PPa, PPe, AGK, AB; Software: AP, MZ, PPe, AGK; Validation: AP, MZ, AB; Formal analysis: AP, MZ, AB; Investigation: AP, MZ, AB; Resources: AB; Data Curation: AP, MZ, JL; Writing - Original Draft: AP; Writing - Review & Editing: AP, MZ, JL, PK, AG, PPa, PPe, AGK, AB; Visualization: AP, AB; Supervision: AB; Project administration: AB; Funding acquisition: AB

## Funding

This research was funded by the National Science Centre, Poland. Research project OPUS-19 (No. Reg. 2020/37/B/NZ9/01499 awarded to AB) and SONATA BIS 10 (No. Reg. 2020/38/E/NZ9/00033 awarded to AB).

## Acknowledgments

We would like to thank Urszula Czech from the University of Agriculture in Poland, Marta Sowa, Katarzyna Pierzchała, Florian Brzostyński and Jacek Matyjaszek for their excellent care of the greenhouse plants. The plasmid kit used to generate plant transformation constructs was a gift from Jan Lohmann (Addgene kit #1000000036).

## Conflict of interest

The authors declare that the research was conducted without any commercial or financial relationships that could be construed as a potential conflict of interest.

## Ethics approval and consent to participate

The plant materials used in this study comply with relevant institutional, national, and international guidelines and legislation. Seeds of *F. tataricum* (sample k-17) are from the N. I. Vavilov Institute of Plant Genetic Resources collections, Saint Petersburg, Russia. The Plant Cytogenetic and Molecular Biology Group Institute of Biology, Biotechnology and Environmental Protection, Faculty of Natural Sciences, University of Silesia in Katowice, Poland, multiplied the obtained seeds. *F. tataricum* sample k-17 is a common cultivated landrace of *F. tataricum*, and seeds are available upon request from the publication’s authors. *F. esculentum* cultivar Panda seeds are commercially available and purchased from the Malopolska Plant Breeding company (Poland).

## Data availability

Data will be made available on request.

---

**Fig. S1:**
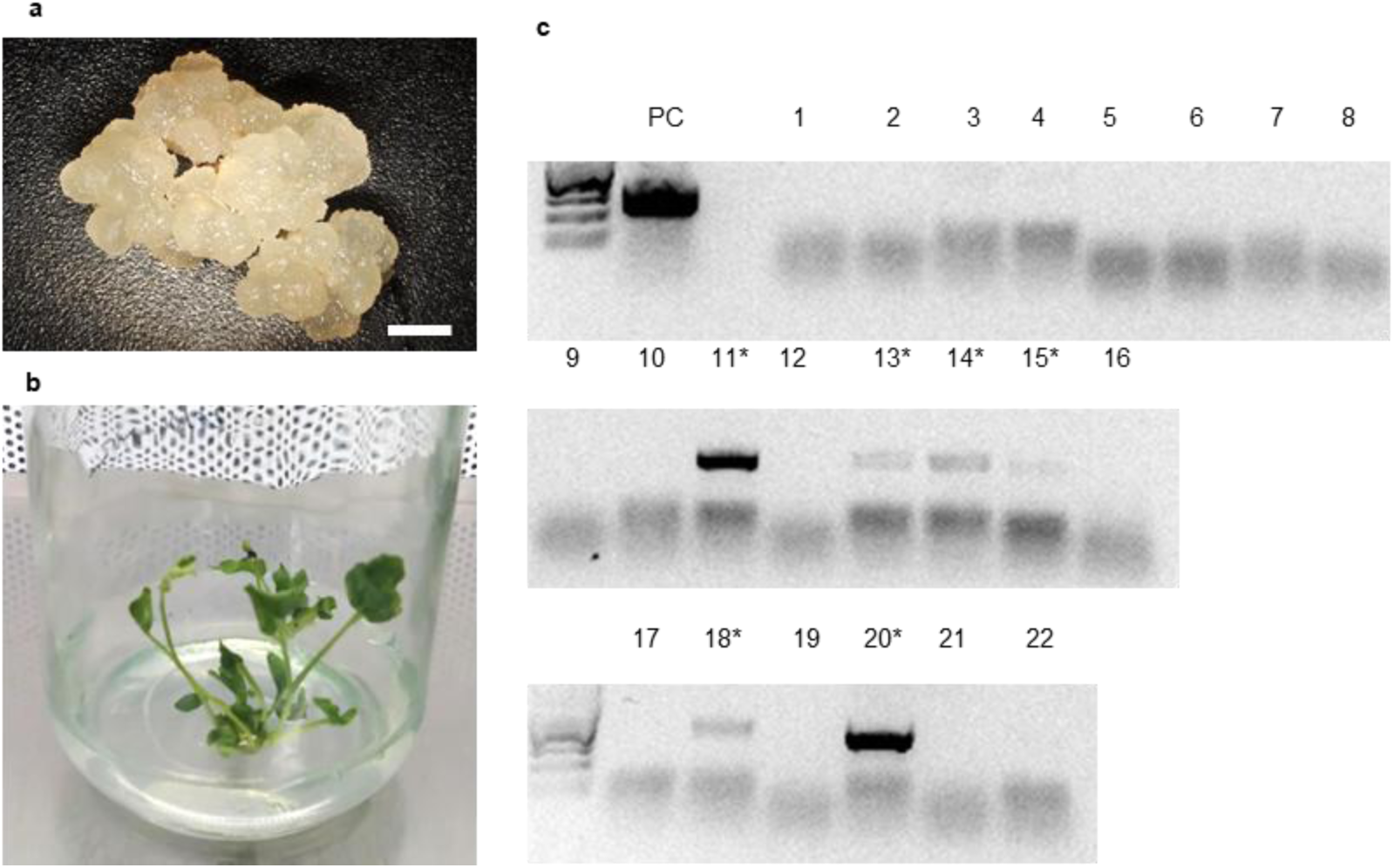
Agrobacterium-mediated transformation of *F. esculentum*. **a**, Callus derived from *F. esculentum* Thrum type. Scale bar, 2 mm. **b**, Regenerated *F. esculentum* plant. **c**, PCR identification of transformed plants with integrated T-DNA (by detection of a fragment of the *Cas9* gene using oligonucleotides Cas9_partial_F and Cas9_partial_R). Plant number 20 carries a mutated *fes-elf3* copy, which was further characterised in this study.

**Fig. S2:**
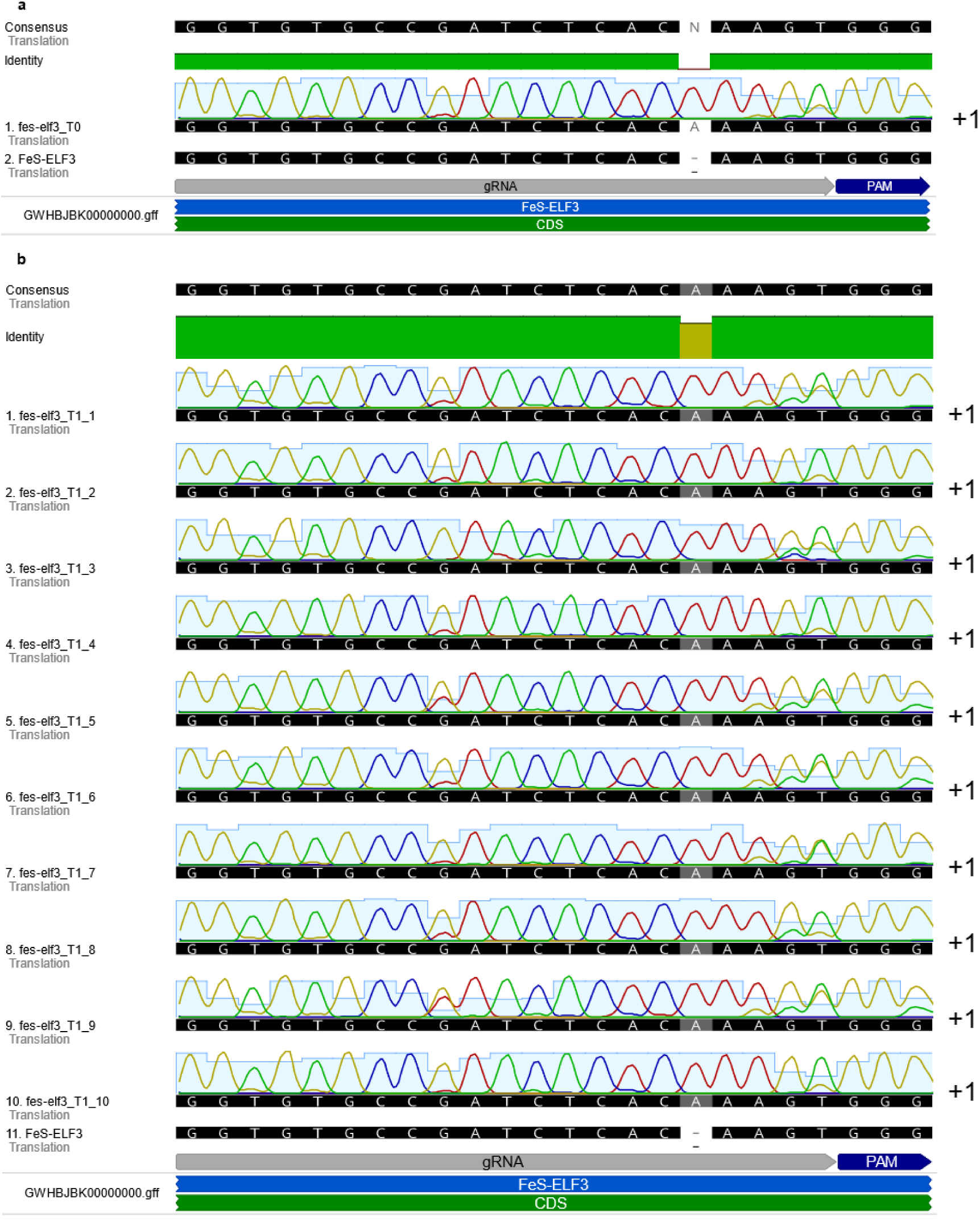
Sequence alignment of *fes-elf3* mutants. (a) Sequence profile of the *fes-elf3* T0 founder line. (b) Representative sequences of ten T1 progeny derived from the *fes-elf3* T0 mutant, demonstrating the stable inheritance of the edited alleles. The WT reference, sgRNA target site, and PAM sequence are indicated.

**Fig. S3:**
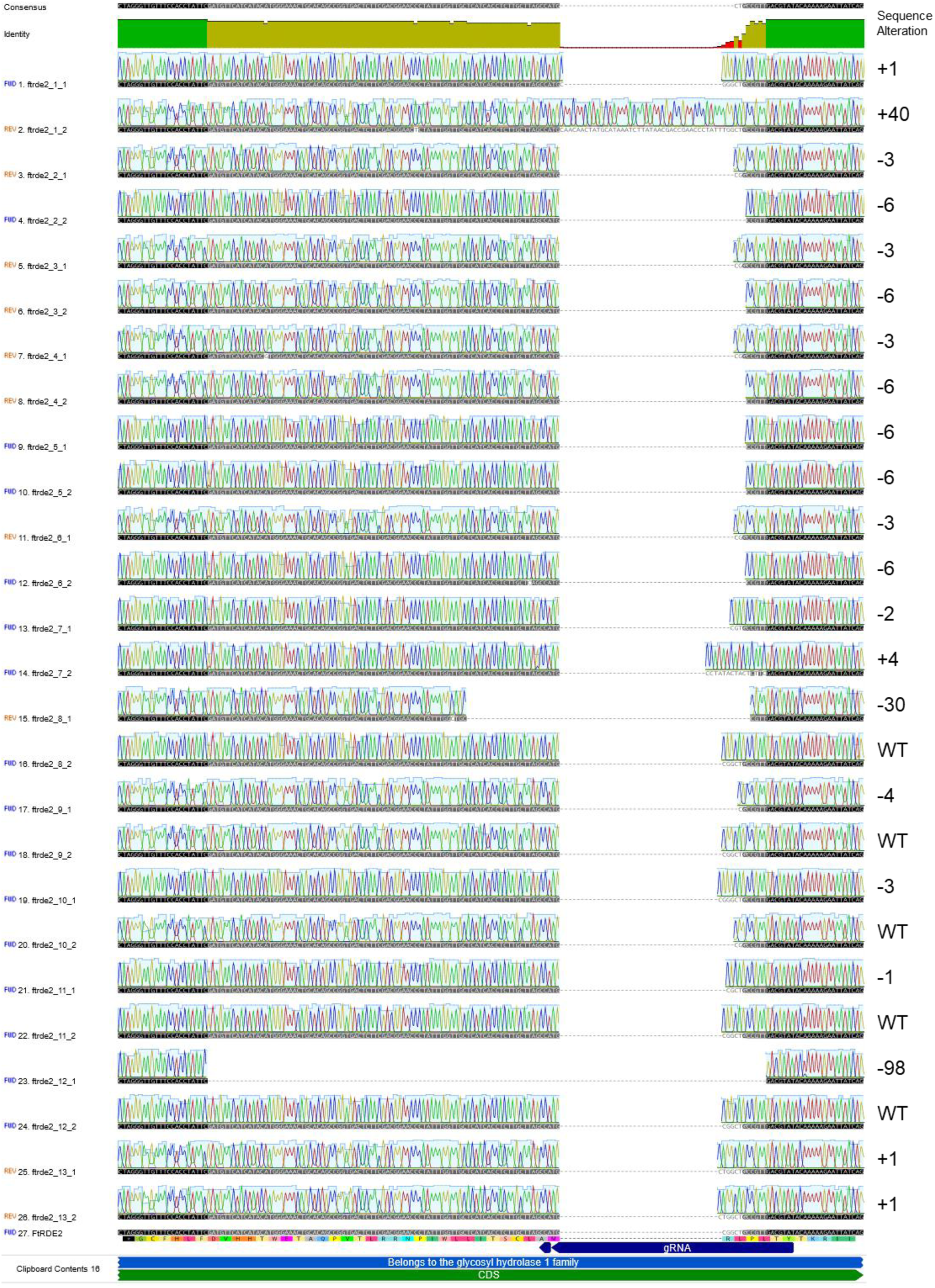
Sequence alignment of the *ftrde2* mutant lines (plants 1–13) in *F. tataricum*. The WT sequence is shown for comparison, with the sgRNA target site and PAM sequence indicated. Various allelic variants are shown, including specific insertions and deletions (indels) resulting from CRISPR/Cas9-mediated editing.

**Fig. S4:**
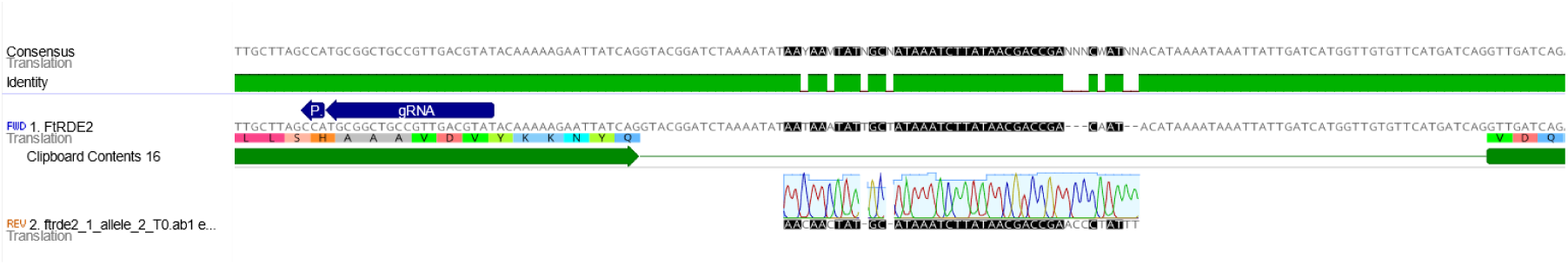
Sequence alignment of the 40-nt insertion in the *ftrde2_1* allele 2 relative to the *FtRDE2* genomic locus.

**Fig. S5:**
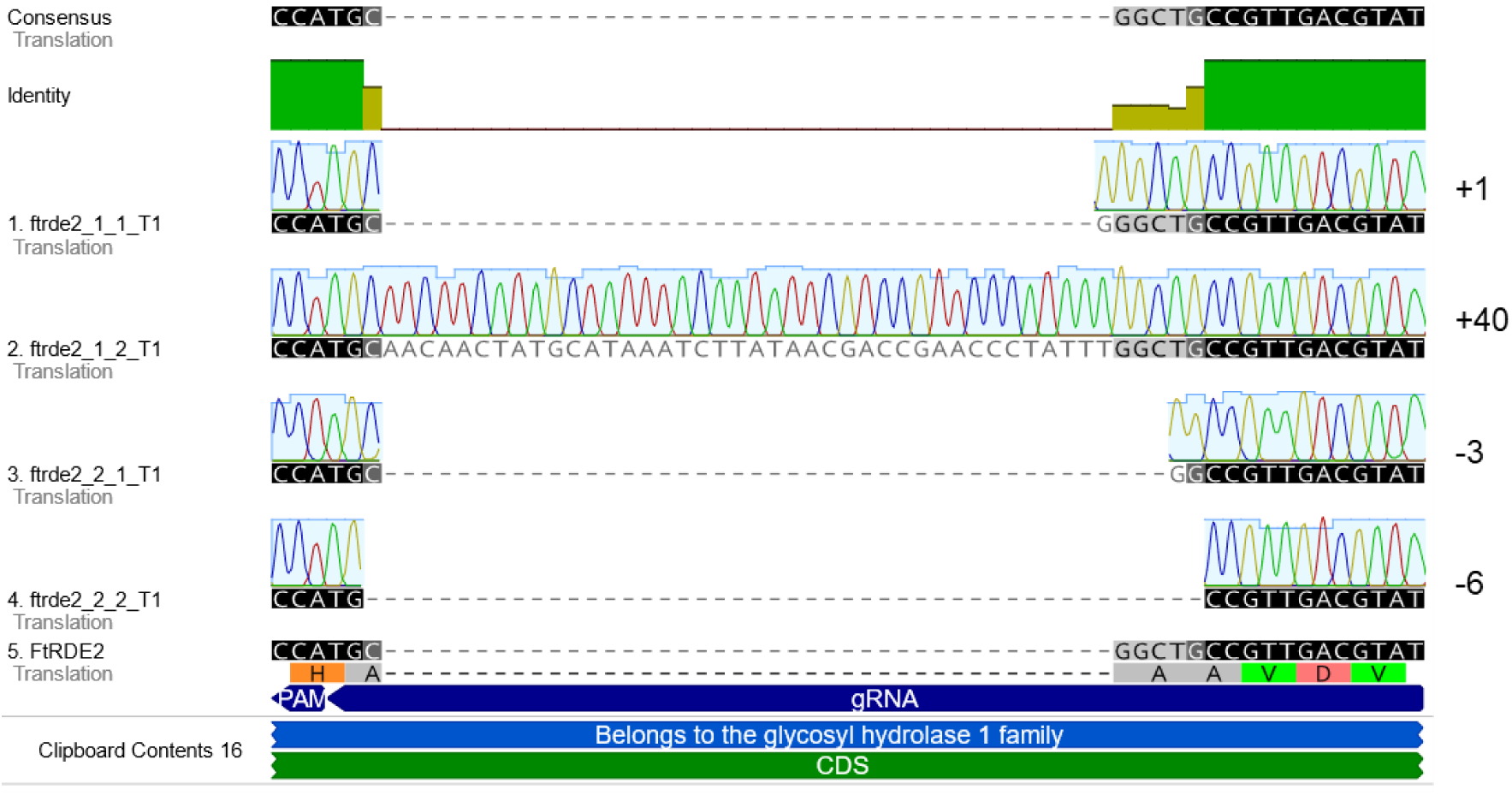
Sequence alignment of T1 progeny from *ftrde2_1* and *ftrde2_2* mutant lines in *F. tataricum*. The WT sequence is provided as a reference, with the sgRNA target site and protospacer adjacent motif (PAM) highlighted. The alignment illustrates the stable inheritance of various allelic variants, including biallelic insertions in *ftrde2_1* and in-frame deletions in *ftrde2_2*.

**Fig. S6.**
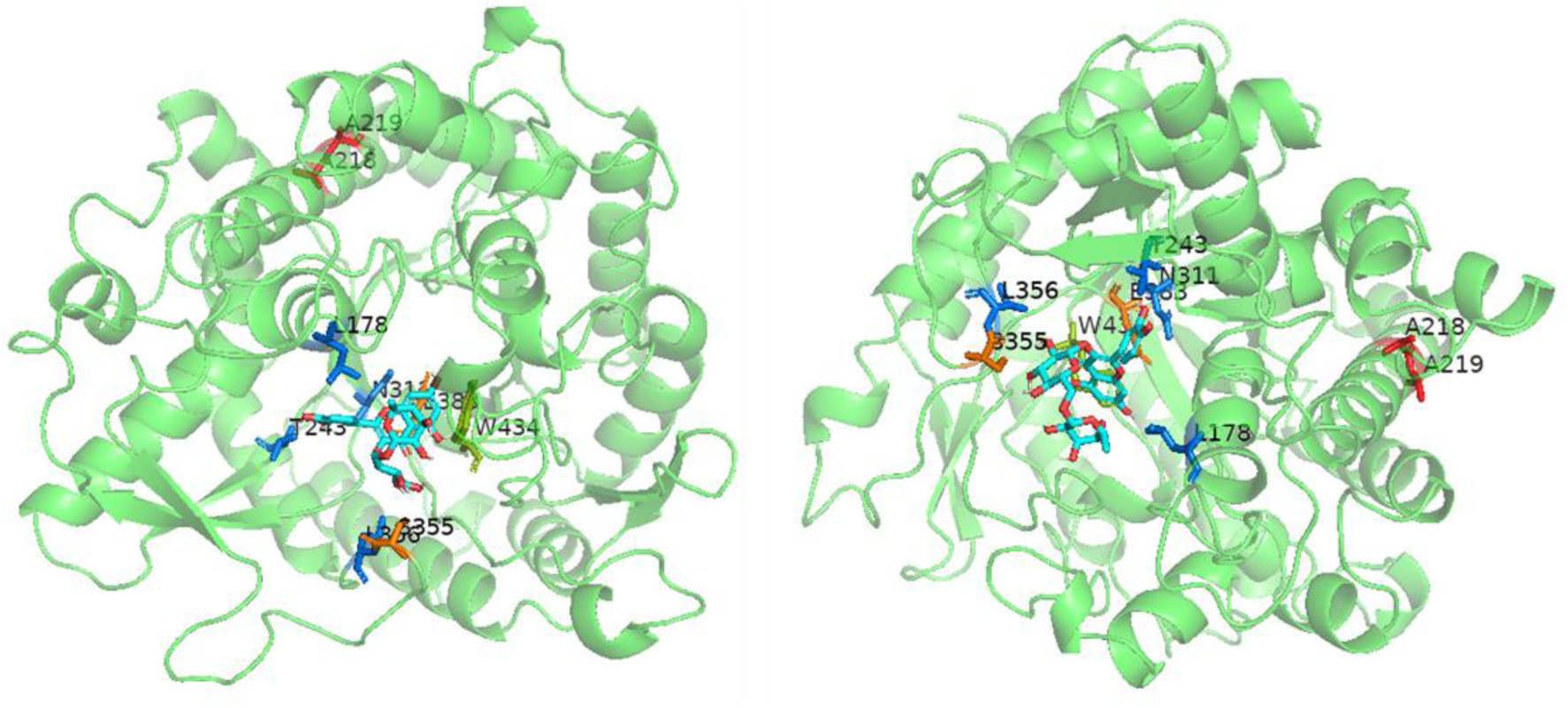
Docking of rutin to the WT FtRDE2 protein. The ligand-binding cavity is formed by LEU-178, ASN-311, GLU-383, LEU-356, and THR-243. In the docked complex, rutin establishes stable hydrogen bonds with GLU-383 and SER-355 and engages in a π–π stacking interaction with TRP-434, supporting a stable binding mode within the active-site region. Residues ALA-218 and ALA-219, which were deleted in the *ftrde2_2* mutant, are highlighted in red. Residue numbering corresponds to the mature protein sequence after removal of the N-terminal signal peptide (residues 1–30), as previously defined (Zhao *et al*., 2025).

**Fig. S7:**
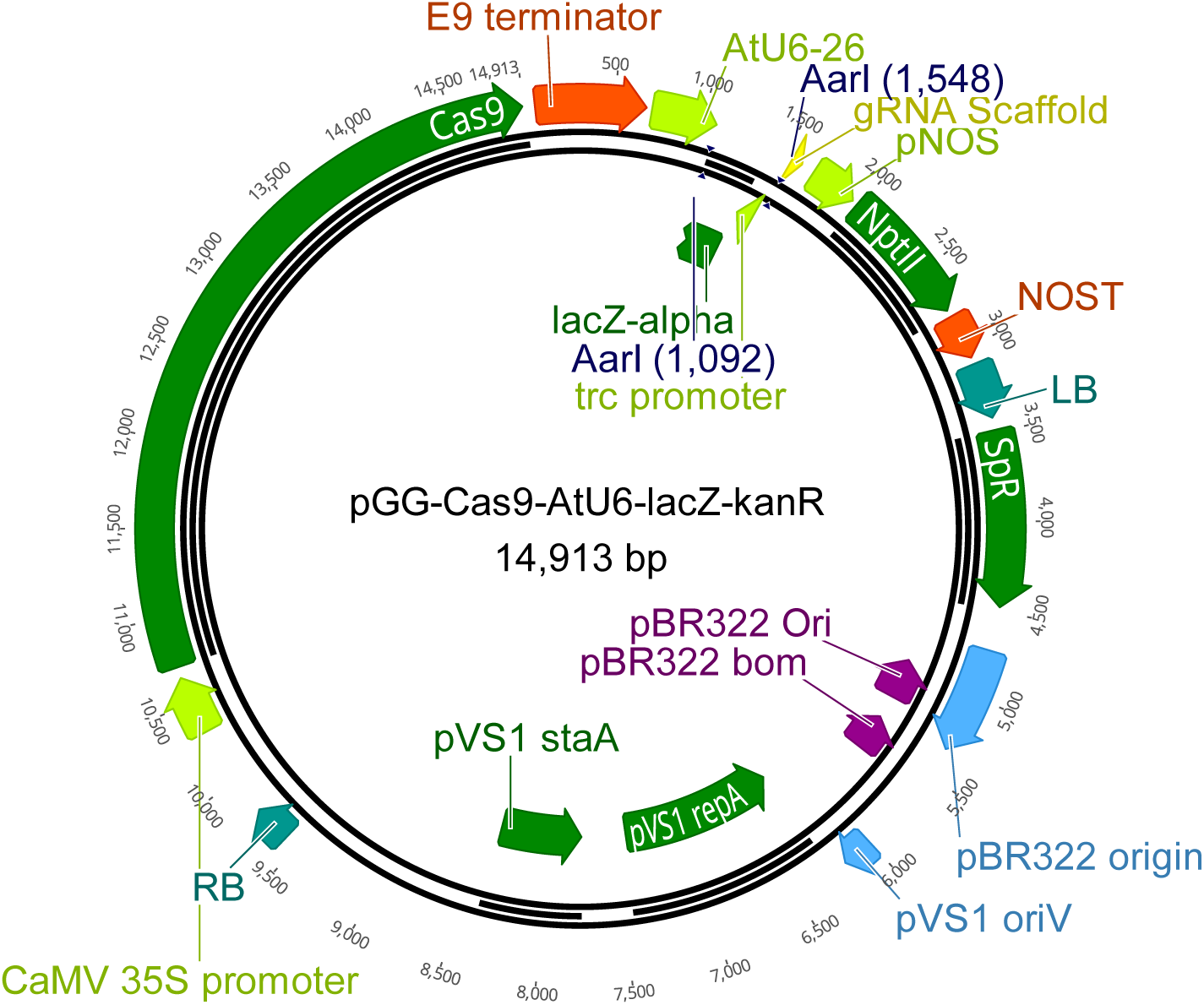
Plasmid map of pGG-Cas9-AtU6-lacZ-kanR, a binary CRISPR/Cas9 vector for sgRNA-mediated targeted mutagenesis in buckwheat. The T-DNA region, flanked by left (LB) and right (RB) borders, contains the AtU6-26 promoter driving sgRNA expression, with the protospacer cloning site defined by flanking AarI restriction sites and the gRNA scaffold. Cas9 is expressed from the CaMV 35S promoter, and the kanamycin plant selectable marker (neomycin phosphotransferase II) from the pNOS promoter. The bacterial backbone carries a spectinomycin resistance cassette (Sp^R^) for bacterial selection, a pBR322 origin, and the pVS1 replicon for stable maintenance in *Agrobacterium tumefaciens*.

**Supplementary Data 1: The Sanger sequencing results of WT sequences and mutants of *F. tataricum* and *F. esculentum*.**

**Supplementary Data 2: Organoleptic analysis of bitterness in *F. tataricum* flour.**

**Supplementary Data 3: AutoDock Vina binding affinity predictions.** Calculated binding affinities (kcal/mol) for rutin docking to FtRDE2 WT, *ftrde2_2_1*(ΔALA218), and *ftrde2_2_1*(ΔALA218ΔALA219) variants. Docking was performed using AutoDock Vina via SwissDrugDesign (https://www.molecular-modelling.ch/swiss-drug-design.html).

| Protein | Model | Calculated affinity (kcal/mol) |
| --- | --- | --- |
| FtRDE2_WT | 1 | -8.499 |
| FtRDE2_WT | 2 | -8.146 |
| FtRDE2_WT | 3 | -7.299 |
| FtRDE2_WT | 4 | -6.034 |
| ***ftRde2_2_1*(ΔALA218)** | **1** | **-8.871** |
| *ftRde2_2_1*(ΔALA218) | 2 | -8.063 |
| *ftRde2_2_1*(ΔALA218) | 3 | -7.992 |
| *ftRde2_2_1*(ΔALA218) | 4 | -7.679 |
| *ftRde2_2_1*(ΔALA218) | 5 | -7.651 |
| *ftRde2_2_1*(ΔALA218) | 6 | -7.303 |
| *ftRde2_2_1*(ΔALA218) | 7 | -6.937 |
| *ftRde2_2_1*(ΔALA218) | 8 | -6.887 |
| *ftRde2_2_1*(ΔALA218) | 9 | -6.871 |
| *ftRde2_2_1*(ΔALA218) | 10 | -6.630 |
| *ftRde2_2_1*(ΔALA218) | 11 | -6.363 |
| *ftRde2_2_1*(ΔALA218) | 12 | -6.358 |
| ***ftRde2_2_2*(ΔALA218ΔALA219)** | **1** | **-8.767** |
| *ftRde2_2_2*(ΔALA218ΔALA219) | 2 | -8.663 |
| *ftRde2_2_2*(ΔALA218ΔALA219) | 3 | -8.142 |
| *ftRde2_2_2*(ΔALA218ΔALA219) | 4 | -7.264 |
| *ftRde2_2_2*(ΔALA218ΔALA219) | 5 | -7.087 |
| *ftRde2_2_2*(ΔALA218ΔALA219) | 6 | -7.058 |
| *ftRde2_2_2*(ΔALA218ΔALA219) | 7 | -6.999 |
| *ftRde2_2_2*(ΔALA218ΔALA219) | 8 | -6.610 |

**Supplementary Data 4: Oligonucleotides used in the study.**

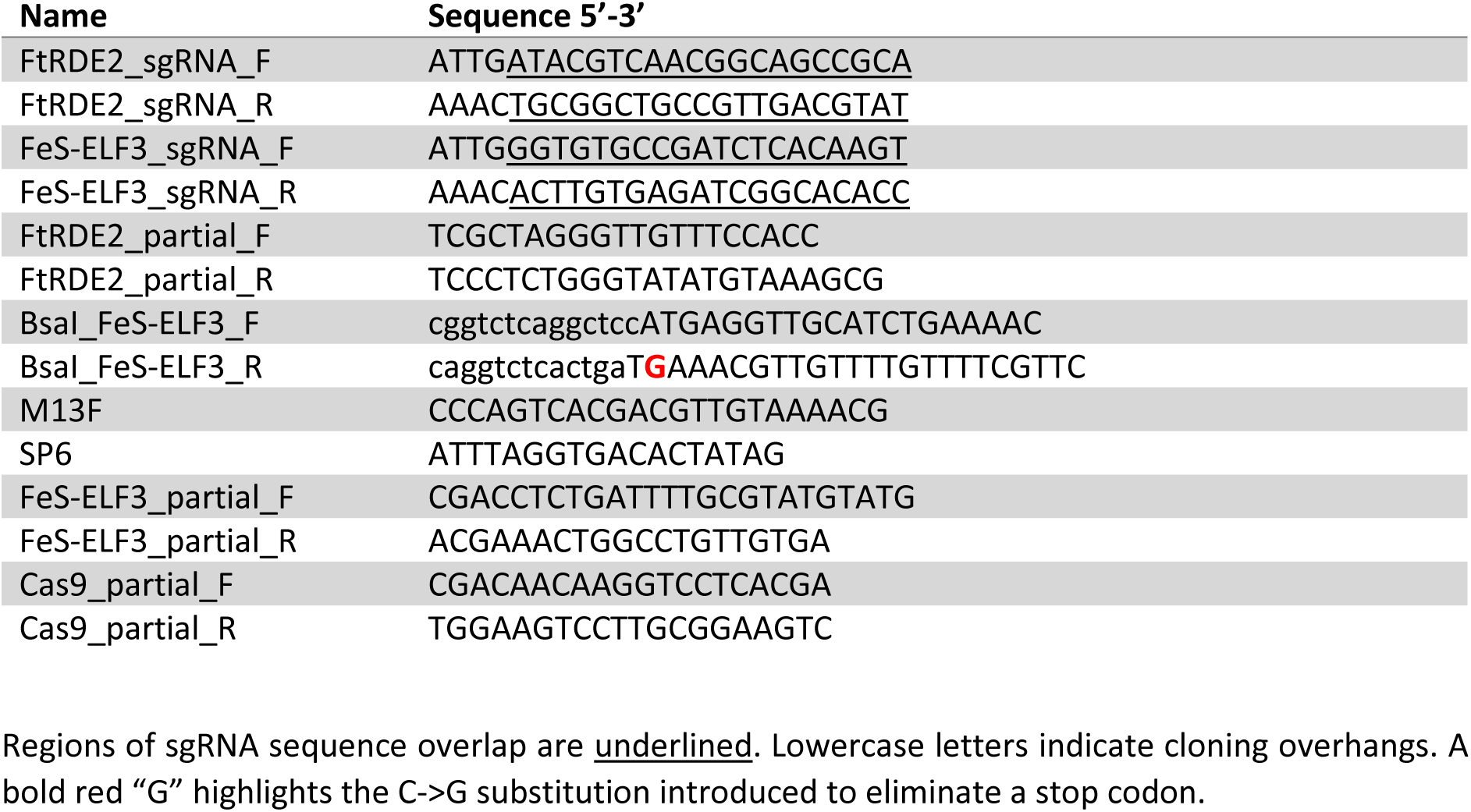

## Notes

### Competing Interest Statement

The authors have declared no competing interest.

